# Observation of reversal in twist-stretch coupling of RNA suggests a unified mechanism for the opposite couplings of DNA and RNA

**DOI:** 10.1101/2021.10.15.464617

**Authors:** Xiao-Wei Qiang, Chen Zhang, Hai-Long Dong, Fu-Jia Tian, Hang Fu, Ya-Jun Yang, Liang Dai, Xing-Hua Zhang, Zhi-Jie Tan

**Affiliations:** Department of Physics and Key Laboratory of Artificial Micro & Nano-structures of Ministry of Education, School of Physics and Technology, Wuhan University, Wuhan 430072, China; College of Life Sciences, the Institute for Advanced Studies, State Key Laboratory of Virology, Hubei Key Laboratory of Cell Homeostasis, Wuhan University, Wuhan 430072, China; Department of Physics, City University of Hong Kong, Hong Kong 999077, China

## Abstract

The functions of DNA and RNA rely on their deformations. When stretched, both DNA and RNA duplexes change their twist angles through twist-stretch coupling. The coupling is negative for DNA but positive for RNA, which is not yet completely understood. Here, our magnetic tweezers experiments show that the coupling of RNA reverses from positive to negative by multivalent cations. Combining with the previously reported tension-induced negative-to-positive coupling-reversal of DNA, we propose a unified mechanism of the couplings of both RNA and DNA based on molecular dynamics simulations. Two deformation pathways are competing when stretched: shrinking the radius causes positive couplings but widening the major groove causes negative couplings. For RNA whose major groove is clamped by multivalent cations and canonical DNA, their radii shrink when stretched, thus exhibiting positive couplings. For elongated DNA whose radius already shrinks to the minimum and canonical RNA, their major grooves are widened when stretched, thus exhibiting negative coupling.

---

Deformations of DNA and RNA play crucial roles in biological processes, such as DNA packaging[1], DNA-protein interactions[2], transcription, translation and material applications such as DNA origami[3]. These deformations can be caused by force[4-6], protein binding [2], ions[7-11], and temperature changes[12,13]. During deformations, many structural parameters of DNA and RNA exhibit couplings, such as twist-stretch coupling[4,14,15] and twist-bend couplin [1,16]. These couplings make significant impacts on mechanical responses of DNA and RNA[4,14,15] and DNA packaging[1]. Experiments observed that stretching a DNA duplex increased DNA twist[4,14] but stretching an RNA duplex decreased RNA twist[15].

The puzzle that DNA and RNA have opposite twist-stretch couplings is striking and has aroused intensive research interests[17,18]. Liebl et al.[17] and Marin-Gonzalez et al.[18] employed state-of-art molecular dynamics simulations to analyze the changes in DNA and RNA structures during stretching and obtained structural parameters affecting the couplings. It was pointed out that the oxygen atom that was present in RNA but absent in DNA obstructed the rotation of the sugar pucker angle, which was a leading reason for the different couplings between RNA and DNA[18]. In this work, we aim to solve the puzzle of the opposite couplings of DNA and RNA by a combination of experiments and simulations and reveal their similarities in nature. Eventually, we want to propose a unified mechanism of twist-stretch couplings of RNA and DNA duplexes.

Here, we employed single-molecule magnetic tweezers (MT) to measure the twist-stretch coupling of RNA and DNA duplexes using torsion-constrained constructs [Fig. 1(a)]. We acquired all data at pH 8.0, 20°C and different salt conditions. The details about the preparation of the DNA and RNA constructs and twist-stretch coupling measurements can be found in the Supplementary Material (Figs. S1-S2) and our previous works[19-21]. We used d*L*/d*N* to characterize twist-stretch coupling[4,22], where d*N* was the change of helical turn and d*L* was the change in extension of the ∼13.7 kbp DNA or RNA duplexes. According to the elastic model[4,22], d*L*/d*N* ∝-g/S, where g is the twist-stretch coupling and S is the stretch modulus with stable positive values [4,15]. Thus, d*L*/d*N* >0 and d*L*/d*N* <0 correspond to the negative and positive coupling, respectively. We measured d*L*/d*N* for DNA duplex as a function of the force also [Fig. 1(c)]. The results reproduced the tension-induced reversal of coupling of DNA reported by Gore et al.[4].

**FIG. 1.**
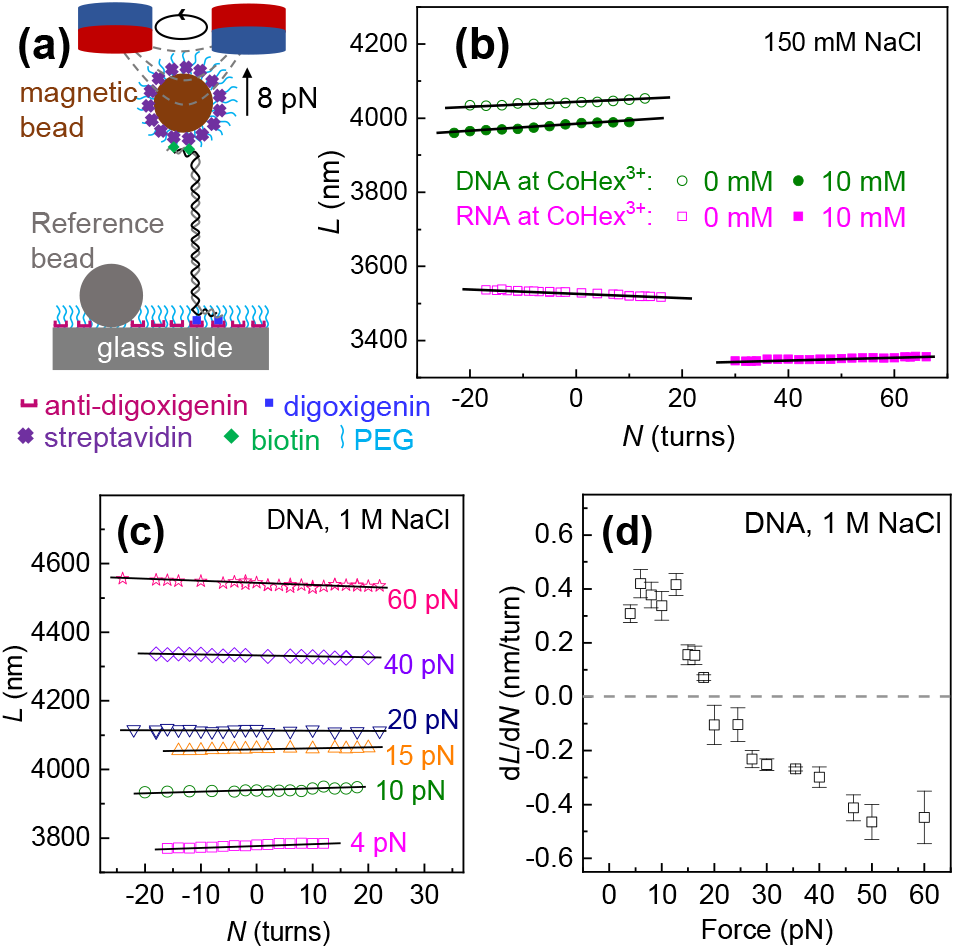
Measuring the twist-stretch couplings using MT. (a) Experimental setup of MT. The torsion-constrained DNA or RNA was anchored between to a superparamagnetic microbead and a glass slide. Then, the bead and glass surfaces were passivated with NHS-mPEG, MW 2K. (b) Representative extensions (*L*) versus rotation turns (*N*) were obtained around the torsional relax point of DNA or RNA at 8 pN using MT. *L* was fitted to linear functions of *N* (black lines). (c) *L* versus *N* for DNA duplex under various stretching forces. (d) d*L*/d*N* as a function of the stretching force for DNA.

Then, we measured the effects of CoHex^3+^ concentration (*C*_Co_) on d*L*/d*N* [Fig. 1(b)] at 150 mM NaCl. For DNA, d*L*/d*N* slightly varied and kept positive (negative g) with the increase of *C*_Co_ as shown in Fig. 2c. For RNA, the positive coupling (d*L*/d*N* <0) at *C*_Co_ = 0 was consistent with previous results [15,17,18,23,24]. However, d*L*/d*N* surprisingly reversed from negative to positive with the increase of *C*_Co_. The reversal occurred at *C*_Co_ ≈0.3 mM. At *C*_Co_ =0 and *C*_Co_ = 100 mM, d*L*/d*N* were ∼ −0.6 and 0.6 nm/turn, respectively. Similarly, we measured the effects of Mg^2+^ concentration (*C*_Mg_) at 0, 10 and 50 mM NaCl. At 10 mM NaCl, Mg^2+^ barely affected the coupling of DNA and d*L*/d*N* kept ∼0.5 nm/turn as shown in Fig. 2b. For RNA duplex, however, with the increase of *C*_Mg_, d*L*/d*N* reversed from ∼ −0.6 nm/turn at *C*_Mg_ = 0 to ∼ 0.3 nm/turn at *C*_Mg_ = 100 mM and then reversed again to d*L*/d*N* to ∼ −0.2 nm/turn at *C*_Mg_ = 2 M. The *C*_Mg_ caused the similar two reversals in d*L*/d*N* at 0 mM NaCl (Fig. S3a). At 50 mM NaCl (Fig. S3b), the *C*_Mg_ affected d*L*/d*N* in the similar trend (increase then decrease for RNA) as the trend at 10 and 0 mM NaCl, whereas *C*_Mg_ did not reverse the coupling of RNA duplex, which might be due to the competitive binding between Mg^2+^ and Na^+^[25]. To summarize, our MT measurements showed that multivalent cations such as CoHex^3+^ and Mg^2+^ of high concentrations reversed the coupling of RNA but not DNA. Our MT experiments reproduced the tension-induced reversal of coupling of DNA as well[4].

**FIG. 2.**
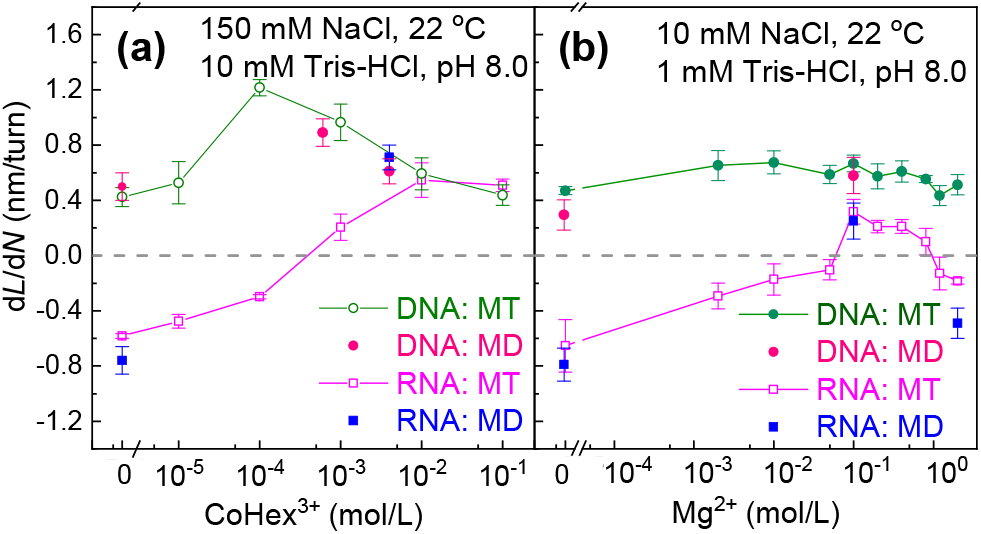
The d*L*/d*N* as functions of CoHex^3+^ (a) and Mg^2+^ (b) concentrations. The experimental d*L*/d*N* obtained from more than four molecules were plotted as data points and error bars. The error bars for the MD data denoted the standard deviations from the values for four equal intervals in equilibrium.

To explore the underlying physical mechanism of effects of multivalent cations on the coupling of DNA and RNA duplexes, we performed all-atom MD simulations with the DNA sequence CGACTCTACGGCATCTGCGC [26] and the same sequence for RNA with T bases replaced by U ones. For simulations, the initial structures of B-DNA and A-RNA were built using the nucleic acid builder of AMBER [19,23,24,27,28]. The force fields of CoHex^3+^, Mg^2+^ and Na^+^ cations were described by the recently proposed ion models [19,29-31]. The bulk cation concentrations were confirmed before the production runs of 1000 or 600 ns[32], and all microscopic structural parameters for DNA and RNA duplexes were calculated using the program Curves+[33]. Please see Figs. S4-6, Table S1 and our previous works[19,23] for the MD simulations, the calculations of the structural/elastic parameters and cation distributions.

As shown in Fig. 2, the calculated d*L*/d*N* from our simulations fairly agreed with experimental results. Notably, our simulations successfully reproduced the reversal of d*L*/d*N* of RNA by CoHex^3+^ (blue squares in Fig. 2a). Furthermore, our simulations also captured the twice reversals of d*L*/d*N* of RNA by Mg^2+^ (blue squares in Fig. 2b). Successful reproduction of the coupling reversals in simulations indicated that our simulations captured the major structural features and interactions that drive the coupling reversals. Thus, we next carefully examined DNA and RNA structural parameters in simulations to unveil the mechanism of the coupling reversals.

Fig. 3 presents the relationships between helical rise h, helical twist ω, helical radius r, major groove width D, and minor groove width d of RNA duplex in the simulations at 0 and 4 mM CoHex^3+^. Several aspects can be inferred from these results. First, the correlation of *h* and ω was reversed by CoHex^3+^ [Fig. 3(a)], which corresponded to the coupling reversal in experiments. Second, in the absence of CoHex^3+^, D strongly correlated with both *h* and *ω* (blue symbols in Fig. 3c-d), which suggested that D mediated the correlation of *h* and *ω*. Third, 4 mM CoHex^3+^ eliminated the correlations of D with *h* and ω, which suggested that D no longer mediated the correlation of *h* and *ω* (orange symbols in Fig. 3c-d). Fourth, in absence of CoHex^3+^, *r* did not correlate obviously with *h* (blue symbols in Fig. 3e), while at 4 mM CoHex^3+^, *r* correlated with *h* [Fig. 3(e)]. Lastly, *r* correlated with *ω* for both in the presence and absence of CoHex^3+^ [Fig. 3(f)]. These results in Fig. 3 agree with previous simulation studies of twist-stretch couplings in the absence of CoHex^3+^ [17,18]. Eventually, we found that RNA radius and major groove width played crucial roles in the coupling reversals, which will be elaborated in further detail below.

**FIG. 3.**
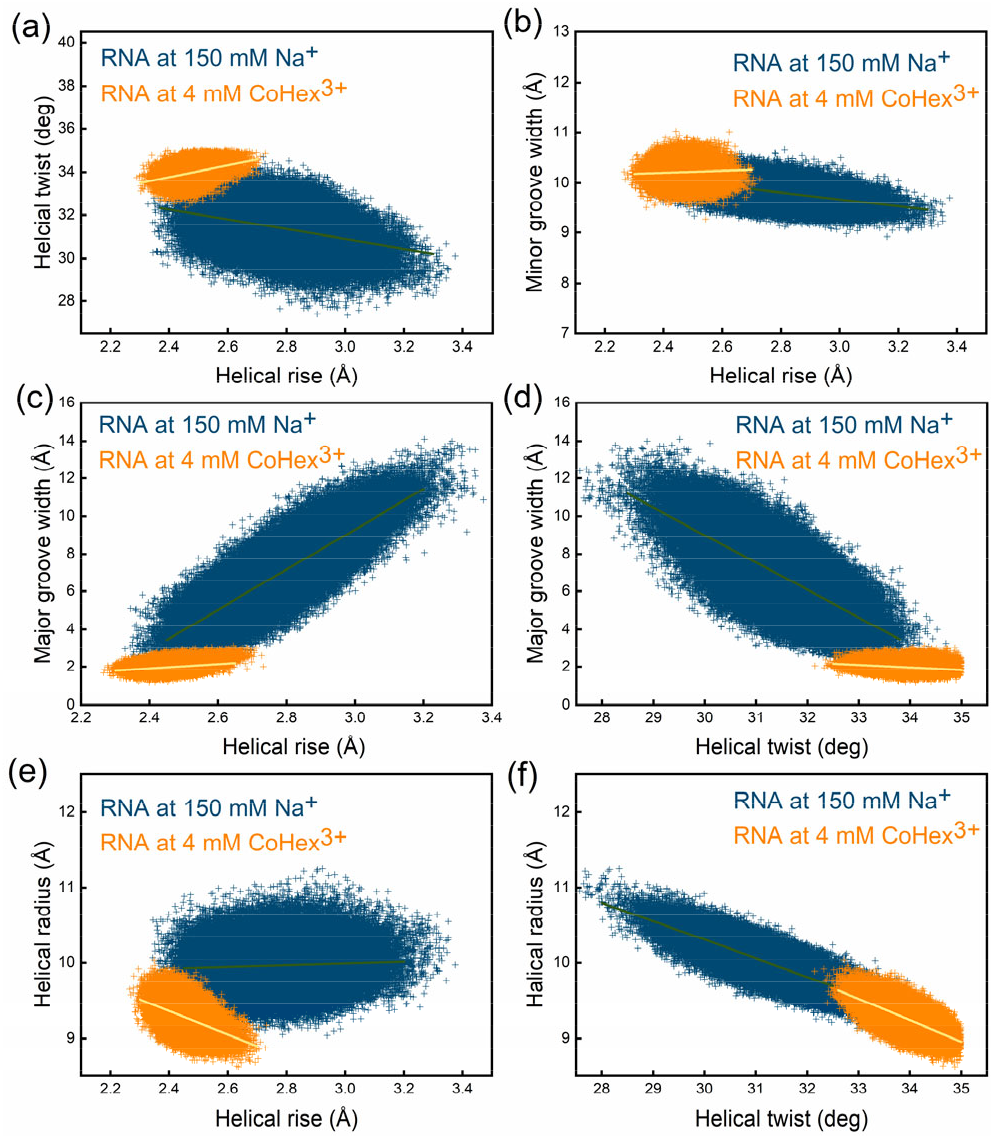
The relationships between helical rise and helical twist from the MD trajectories. Error bars showed standard errors of the averages in each interval. In each panel, the raw data were shown as a density landscape with normalized probability.

The results in Fig. 3 revealed the importance of the major groove width in the reversal of twist-stretch coupling of RNA. To further elaborate on the role of the major groove width, we carried out the following analysis. We calculated average major groove width ⟨*D*⟩ and its fluctuation *σ*_*D*_ and obtained 6.9 ± 1.7 and 2.0 ± 0.2 Å for 0 and 4 mM CoHex^3+^. The dramatic decreases in ⟨*D*⟩ and *σ*_*D*_ indicated the narrowing and clamping by CoHex^3+^, respectively. The narrowing and clamping effects were attributed to the most preferred binding of CoHex^3+^ in the deep major groove of the RNA duplex. The preferences of binding sites depended on the surface electrostatic potentials calculated with the APBS as shown in Fig. 4a-b [34]. Note that DNA had a different surface electrostatic potential and CoHex^3+^ preferred to bind on the DNA phosphate backbone (Table S2). Accordingly, for DNA, there was no narrowing and clamping of the major groove by CoHex^3+^. For RNA, Mg^2+^ had a weaker clamping effect than CoHex^3+^ due to its lower ion charge. Thus, a significant clamping effect required higher *C*_Mg_. An intriguing phenomenon was that when *C*_Mg_ exceeded a threshold, the clamping effect became even weaker. The reason was as follows. For high *C*_Mg_, e.g., 2 mol/L, RNA major grooves were fully occupied by Mg^2+^, and the excess Mg^2+^ bound to RNA phosphate backbone, which weakened the clamping effect because phosphate groups experienced attractions from Mg^2+^ both in major grooves and on phosphate groups (Table S2). The weakening of the clamping effect at very high *C*_Mg_ should be responsible for the second reversal of twist-stretch coupling of Fig. 2b.

**FIG. 4.**
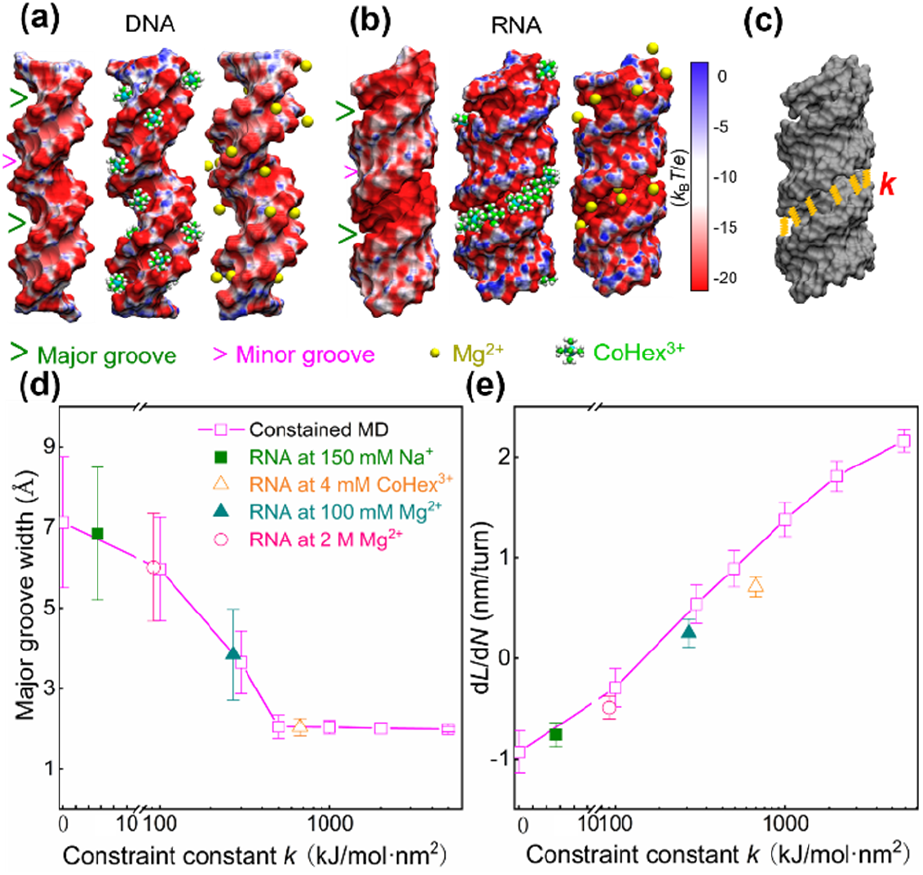
The effects of major groove clamping by multivalent cations. (a, b) Representative structures of DNA and RNA showing surface electrostatic potentials and the binding of CoHex^3+^ were revealed by MD simulations. In each panel, left: standard form duplexes; middle: in CoHex^3+^ solutions; right: in Mg^2+^ solutions. (c) An illustration for the RNA duplex with harmonic constraint between phosphorus atoms across the major groove used for our MD simulations, and the spring constant is k. (d) The relationship between D and k for the RNA duplex, and the error bars denoting the standard deviations. (e) d*L*/d*N* of the RNA duplex as a function of k. The k values for typical cation conditions corresponded to those in panel (e).

To confirm that CoHex^3+^ reverses twist-stretching coupling through narrowing and clamping major groove, we performed additional MD simulations for RNA duplex with artificial restraints of major groove widths in the absence of CoHex^3+^. These artificial restraints were imposed through springs that tend to bring together phosphorus atoms across the major groove, as illustrated in Fig. 4c (see more details in Supplementary Methods and Fig. S7). As we increased the spring constant, the major groove width and its fluctuation decreased [Fig. 4(d)], and simultaneously, d*L*/d*N* reversed from negative to positive values [Fig. 4(e)], which demonstrated the critical role of narrowing and clamping major groove in the coupling reversal. Furthermore, these constrained simulations also reproduced the increases of bending persistence length and stretch modulus of RNA duplex by CoHex^3+^ (Table S4)[19,35].

Intriguingly, RNA and DNA both can reverse their twist-stretch coupling. For canonical DNA, the deformation upon stretching shrinks its radius, which causes overwinding, as proposed by Gore et al[4]. The situation resembles the RNA with CoHex^3+^. For an elongated DNA under a strong stretching force, the DNA radius already reaches its minimum and cannot shrink anymore (Fig. S8). In this case, the DNA resorts to the other deformation pathway, i.e., widening the major groove width, in the response to stretching (Fig. S8). The situation resembles the canonical RNA.

Now we are clear that the sign of twist-stretch coupling depends on the deformation pathway. The next question is what determines the deformation pathway in DNA and RNA during stretching. Naturally, RNA or DNA adopts the deformation pathway with the lowest energy during stretching. For canonical RNA, the radius is relatively difficult to shrink probably due to the extra oxygen that RNA contains and DNA does not [18]. For RNA at 4 mM CoHex^3+^, we find that narrowing and clamping major groove width by CoHex^3+^ impose a high energy penalty for widening major groove, and hence RNA resorts to the other deformation pathway for stretching. In this situation, radius of RNA shrinking becomes energetically favorable. For canonical DNA, radius shrinking is an energetically favorable deformation pathway[6]. However, the radius cannot decrease to zero due to the finite volumes filled by DNA atoms. Once the minimum radius is reached, DNA must resort to the other deformation pathway.

Apparently, the lengths of DNA and RNA axes play an important role in determining the deformation pathway and the sign of twist-stretch coupling. For relatively short axial lengths in canonical DNA or compressed RNA (Fig. 5(a2)), the radius-shrinking pathway dominates, while for relatively large axial lengths in elongated DNA or canonical RNA, the radius-shrinking pathway is obstructed.

**FIG. 5.**
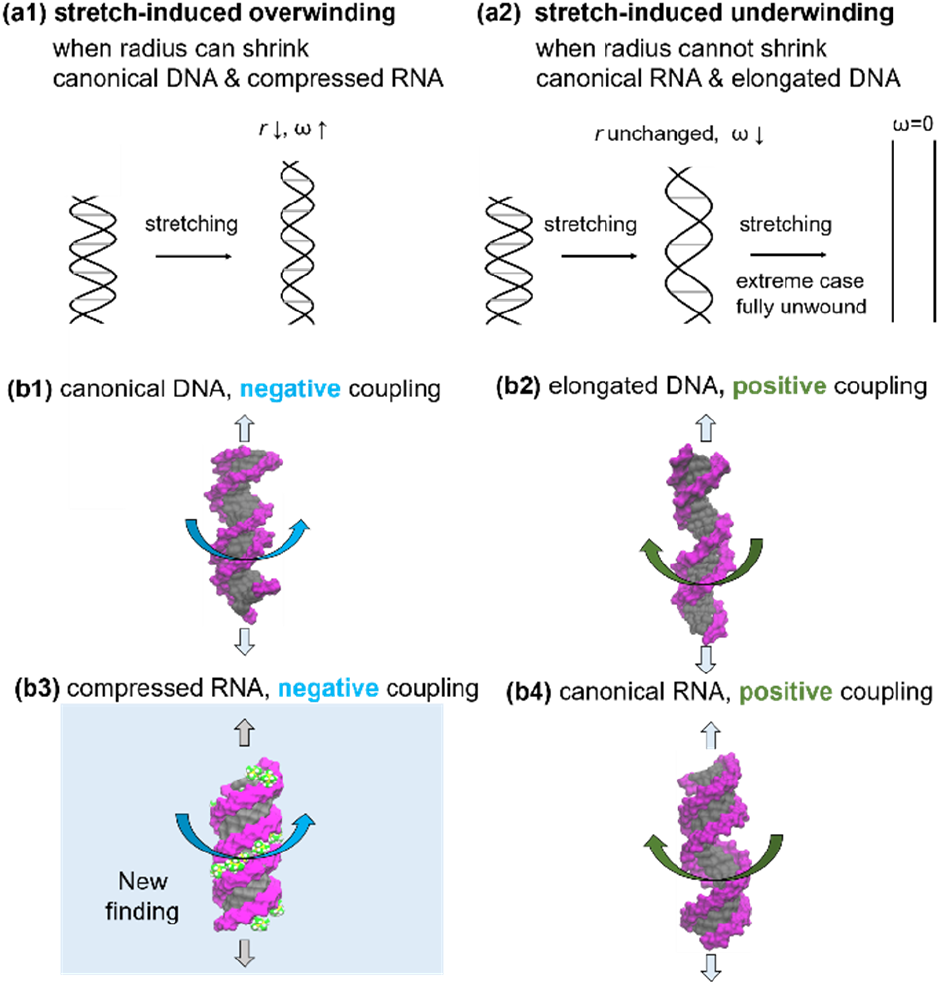
The unified mechanism of twist-stretch couplings of both DNA and RNA. (a) Two deformation pathways for stretch-induced overwinding and underwinding, respectively. (b) Twist-stretch couplings of DNA and RNA duplex.

It is insightful to discuss the extreme case of the full stretch as illustrated by Fig. 5(a2). When the full stretch is achieved, the twist is zero. This extreme case indicates that stretch-induced underwinding always occurs for sufficient strong forces or sufficiently elongated DNA and RNA.

It is worth pointing out that two deformation pathways are not mutually exclusive. It is quite likely that in the intermediate regime, both deformation pathways take place. Accordingly, stretch-induced twist change is contributed by both deformation pathways, and the observed twist change is the net effect. In particular, shrinking of the radius by itself does not directly contribute to stretch. It means that even when the radius shrinking dominates, there are still substantial deformations in minor and major grooves. As shown by Fig. 3(b), the stretching of the RNA with CoHex^3+^ induces the slight widening of minor grooves beside the shrinking of radius.

The reason why the radius shrinking is very important in twist-stretch coupling is that twist-radius correlation is prevalent and strong[18]. Recall that twist-radius correlation takes effect for both 0 and 4 mM CoHex^3+^ as shown in Fig. 4(f). The radius-shrinking deformation pathway significantly contributes to overwinding, which may overwhelm underwinding induced by other deformation pathways.

In summary, we revealed a unified mechanism of twist-stretch couplings of RNA and DNA duplexes. Our MT experiments and MD simulations revealed that multivalent cations reversed the twist-stretch coupling of RNA duplex, which filled in the last piece [Fig. 5(b)] and completed the picture of twist-stretch couplings of RNA and DNA duplexes. Thus, both DNA and RNA can exhibit positive and negative twist-stretch couplings depending on the dominant deformation pathways during stretching. If the radius-shrinking pathway dominates, stretch induces overwinding. If the widening of the major groove dominates, stretch induces underwinding. Which deformation pathway dominates depends on the states of DNA and RNA structures at the time when stretched and also depends on the structural restraints. Narrowing and clamping RNA major groove width obstructs the deformation through widening the major groove, thus RNA adopts the radius-shrinking pathway during stretching, which leads to stretch-induced overwinding. Our results enrich the understanding of mechanical responses of DNA and RNA, which may be useful in rational control of RNA and DNA structures in material applications[3,36].

We are grateful for the financial support from the National Natural Science Foundation of China (Nos. 11774272, 11704286, 12075171 and 31670760), valuable discussions with Profs. Shi-Jie Chen and Xiangyun Qiu, and the Super Computing Center of Wuhan University.

## Supporting information

Supplementary simulation method, Figures 1-8, Tables 1-3

## References

[1] E. Skoruppa, S. K. Nomidis, J. F. Marko, and E. Carlon, Phys. Rev. Lett. 121, 088101 (2018).

[2] S. Kim et al., Science 339, 816 (2013).

[3] H. Dietz, S. M. Douglas, and W. M. Shih, Science 325, 725 (2009).

[4] J. Gore, Z. Bryant, M. Nollmann, M. U. Le, N. R. Cozzarelli, and C. Bustamante, Nature 442, 836 (2006).

[5] P. Gross, N. Laurens, L. B. Oddershede, U. Bockelmann, E. J. G. Peterman, and G. J. L. Wuite, Nat. Phys. 7, 731 (2011).

[6] Z. Bryant, M. D. Stone, J. Gore, S. B. Smith, N. R. Cozzarelli, and C. Bustamante, Natur 424, 338 (2003).

[7] P. Anderson and W. Bauer, Biochemistry 17, 594 (1978).

[8] F. Kriegel, N. Ermann, R. Forbes, D. Dulin, N. H. Dekker, and J. Lipfert, Nucleic Acids Res. 45, 5920 (2017).

[9] S. J. Chen, Annu.Rev. Biophys. 37, 197 (2008).

[10] L. Z. Sun, D. Zhang, and S. J. Chen, Annu.Rev. Biophys. 46, 227 (2017).

[11] I. S. Tolokh, S. A. Pabit, A. M. Katz, Y. Chen, A. Drozdetski, N. Baker, L. Pollack, and A. V. Onufriev, Nucleic Acids Res. 42, 10823 (2014).

[12] F. Kriegel, C. Matek, T. Drsata, K. Kulenkampff, S. Tschirpke, M. Zacharias, F. Lankas, and J. Lipfert, Nucleic Acids Res 46, 7998 (2018).

[13] D. E. Depew and J. C. Wang, Proc. Natl. Acad. Sci. U.S.A. 72, 4275 (1975).

[14] T. Lionnet, S. Joubaud, R. Lavery, D. Bensimon, and V. Croquette, Phys. Rev. Lett. 96, 178102 (2006).

[15] J. Lipfert et al., Proc. Natl. Acad. Sci. U.S.A. 111, 15408 (2014).

[16] S. K. Nomidis, F. Kriegel, W. Vanderlinden, J. Lipfert, and E. Carlon, Phys. Rev. Lett. 118, 217801 (2017).

[17] K. Liebl, T. Drsata, F. Lankas, J. Lipfert, and M. Zacharias, Nucleic Acids Res. 43, 10143 (2015).

[18] A. Marin-Gonzalez, J. G. Vilhena, R. Perez, and F. Moreno-Herrero, Proc. Natl. Acad. Sci. U.S.A. 114, 7049 (2017).

[19] H. Fu, C. Zhang, X. W. Qiang, Y. J. Yang, L. Dai, Z. J. Tan, and X. H. Zhang, Phys. Rev. Lett. 124, 058101 (2020).

[20] Y. J. Yang et al., JACS 142, 9203 (2020).

[21] C. Zhang, H. Fu, Y. Yang, E. Zhou, Z. Tan, H. You, and X. Zhang, Biophys. J. 116, 196 (2019).

[22] J. Wei, L. Czapla, M. A. Grosner, D. Swigon, and W. K. Olson, Proc. Natl. Acad. Sci. U.S.A. 111, 16742 (2014).

[23] L. Bao, X. Zhang, Y. Z. Shi, Y. Y. Wu, and Z. J. Tan, Biophys. J. 112, 1094 (2017).

[24] J. H. Liu, K. Xi, X. Zhang, L. Bao, X. Zhang, and Z. J. Tan, Biophys. J. 117, 74 (2019).

[25] K. Xi, F. H. Wang, G. Xiong, Z. L. Zhang, and Z. J. Tan, Biophys. J. 114, 1776 (2018).

[26] R. S. Mathew-Fenn, R. Das, and P. A. Harbury, Science 322, 446 (2008).

[27] D. A. Case et al., J. Comput. Chem. 26, 1668 (2005).

[28] A. Srivastava, R. Timsina, S. Heo, S. W. Dewage, S. Kirmizialtin, and X. Qiu, Nucleic Acids Res. 48, 7018 (2020).

[29] P. F. Li, B. P. Roberts, D. K. Chakravorty, and K. M. Merz, J. Chem. Theory Comput. 9, 2733 (2013).

[30] T. Sun, A. Mirzoev, N. Korolev, A. P. Lyubartsev, and L. Nordenskiöld, J. Phys. Chem. B 121, 7761 (2017).

[31] I. S. Joung and T. E. Cheatham III, J. Phys. Chem. B 112, 9020 (2008).

[32] Y. Y. Wu, Z. L. Zhang, J. S. Zhang, X. L. Zhu, and Z. J. Tan, Nucleic Acids Res. 43, 6156 (2015).

[33] R. Lavery, M. Moakher, J. Maddocks, D. Petkeviciute, and K. Zakrzewska, Nucleic Acids Res. 37, 5917 (2009).

[34] N. A. Baker, D. Sept, S. Joseph, M. J. Holst, and J. A. McCammon, Proc. Natl. Acad. Sci. U.S.A. 98, 10037 (2001).

[35] A. V. Drozdetski, I. S. Tolokh, L. Pollack, N. Baker, and A. V. Onufriev, Phys. Rev. Lett. 117, 028101 (2016).

[36] A. M. Maier, W. Bae, D. Schiffels, J. F. Emmerig, M. Schiff, and T. Liedl, ACS Nano 11, 1301 (2017).

