## Supplementary simulation method, Figures 1-8, Tables 1-3 for "Observation of reversal in twist-stretch coupling of RNA suggests a unified mechanism for the opposite couplings of DNA and RNA"

#### All-atom MD simulations

We used the 20-bp DNA duplex with a sequence of CGACTCTACGGCATCTGCGC and its complementary one according to the experiments on flexibility [4] and the 20-bp RNA duplex with the same as the DNA sequences except that T was replaced by U, the DNA and RNA duplexes are with a CG content of 57%. The initial structures of the DNA duplex in B-form and RNA duplex in A-form were built using the Nucleic Acid Builder of AMBER [5].

The initial structures of DNA and RNA duplexes were immersed in periodic simulation rectangle boxes of 12×12×10 $nm^3$ containing explicit water molecules and cations. The counterions of $Na^+$ and the salt ions of 150 mM NaCl were added with the ion model from Joung and Cheatham [6] and 4 mM $CoHex^{3+}$ cations were added with the ion model from Ref. [7] and Merz/Li ion model for 100 mM $Mg^{2+}$ [8]. To obtain the desirable bulk concentration for mixed $Na^+$/$CoHex^{3+}$ and $Na^+$/$Mg^{2+}$ solutions, following Refs. [9, 10], the simplified implicit-solvent Monte Carlo simulations and the TBI-based calculations were employed to estimate the numbers of $Na^+$/$CoHex^{3+}$ and $Na^+$/$Mg^{2+}$ cations in the simulational boxes before the time-consuming all-atom simulations and the obtained numbers of $CoHex^{3+}$, $Mg^{2+}$ and $Na^+$ cations were used in all simulations, see Refs. [9, 11]. Additionally, to obtain low $Na^+$ concentration (10 mM), a large box (15×15×12 $nm^3$) was also used for both DNA and RNA duplexes in our simulations.

Following the initial setup of the molecular system we employed 5000 steps of energy minimizations to remove the bad contacts that may arise due to random placement of water and cations. All MD simulations were performed in the isothermic-isobaric ensemble (P=1 atm, T=298 K) when the systems were energy-minimized, thermalized and equilibrated. Afterward, 0.6 and 1 microsecond MD simulations were performed for the equilibrated systems at constant temperature (298.15 K) using the Berendsen thermostat, and pressure (1 atm) using Parrinello-Rahman barostat. Periodic boundary conditions were implemented in all directions and the Particle Mesh Ewald method for long-range interactions [12] with a grid spacing of 0.16 nm and an interpolation of order 4. We used a cutoff radius of 1 nm for neighbor search. An integration step of 2 fs was used in conjunction with the leap-frog algorithm [13]. During this process, DNA and RNA duplexes were completely free in the solutions. Covalent bonds of the water and DNA/RNA duplexes were constrained to their equilibrium geometries using LINCS algorithms. All of our simulations were carried out using the Gromacs 4.6 software package with newly refined AMBER ff99bsc1 +$\chi$OL3 force fields and TIP3P water model for molecular interactions. The MD simulations were performed until 1000ns for the cation conditions with $CoHex^{3+}$ and until 600ns for other cation conditions. The MD simulations are nearly converged after ~200 ns for DNA and RNA duplexes with $Na^+$ and $Mg^{2+}$ and converged after ~400 ns for DNA and RNA duplexes with $CoHex^{3+}$ as shown in Fig. S6 and Fig. S7.

#### All-atom MD simulations for RNA duplex with harmonic distance constraints across the major groove

To verify the effect of multivalent cation-clamping across major groove on the twist-stretch coupling of RNA duplex, we performed the constrained all-atom MD simulations for the RNA duplexes with different distance constraints across the major groove. In the simulations, we used the equilibrium structure of RNA in 4 mM $CoHex^{3+}$ as the initial structure and we removed all the $CoHex^{3+}$ cations and added harmonic distance constraints between adjacent phosphate atoms across the major groove; see Fig. 4(d). The harmonic constraint potential between phosphorus atoms across the major groove is given by $v(r_{ij})=k(r_{ij}-r_1)^2/2$ for $r_{ij}<r_2$ [14, 15], where $r_1$ and $r_2$ represent the equilibrium distance and the upper bound for the potential, $r_{ij}$ is the distance between two phosphate atoms across the major groove, and $k$ is the distance constraint constant. In our constrained MD simulations, we set $r_1$=0.8 nm and $r_2$=3 nm in the topology file for the force field, where 0.8 nm is the mean distance between adjacent phosphate atoms across major groove from the above MD simulations with 4mM $CoHex^{3+}$. Extensive constraint constants were used: $k$=0 kJ/mol·nm$^2$, $k$=100 kJ/mol·nm$^2$, $k$=300 kJ/mol·nm$^2$, $k$=500 kJ/mol·nm$^2$, $k$=1000 kJ/mol·nm$^2$, $k$=2000 kJ/mol·nm$^2$, and $k$=5000 kJ/mol·nm$^2$. The constrained MD simulations were performed until 250ns for each covered constraint constant $k$. As shown in Fig. S5, the constrained MD simulations can reach their equilibrium very quickly, and we made the analyses for the trajectories of the last 150 ns.

**Helical parameters of DNA and RNA duplexes**

In this work, the helical/local parameters were all obtained using the program Curves+ [16, 17]. To avoid the end effects of short duplexes, the three base pairs at each end were removed for all the analyses of the MD trajectories [18]. We used the central 14-bp segment taken from the 20-bp DNA and RNA duplexes to analyze the relative flexibility of DNA and RNA duplexes in twist-stretch coupling. It is noted that the CG content is ~57% for the central DNA and RNA duplexes. The calculated helical parameters of DNA and RNA duplexes at typical cation conditions were listed in Table S1 and Fig. S8, including helical rise, helical twist, helical radius, major groove width, minor groove width, and their standard deviations.

**Calculating elastic parameters of DNA and RNA duplexes**

Macroscopic elastic parameters can be evaluated as the diagonal terms of the elastic matrix $K$ associated with the global helical coordinates (contour length $L$ and cumulative helical twist $\Phi$ over the central 14 bp), and are determined by [19, 20]

$$\begin{aligned} K &= k_B T L_0 V^{-1} \\ &= k_B T L_0 \begin{pmatrix} \langle \Delta L^2 \rangle & \langle \Delta L \times \Delta\Phi \rangle \\ \langle \Delta L \times \Delta\Phi \rangle & \langle \Delta\Phi^2 \rangle \end{pmatrix}^{-1} , \end{aligned} \quad \text{(S1)}$$

where $V$ is a 2×2 covariance matrix of the two global helical coordinates ($L$ and $\Phi$) and can be obtained from the MD simulation trajectories [19, 20], $L_0$ is the average value of contour length, $T$ is the thermodynamic temperature in Kelvin, and $k_B$ is the Boltzmann constant. In Eq. S1, the diagonal terms of $V^{-1}$ can be understood as the reciprocal of the partial variances, as described in Noy and Golestanian [20]. The twist-stretch coupling is associated with contour length $L$ and cumulative helical twist $\Phi$ in Eq. S1, respectively. Stiffness analyses are made for the global properties of the central 14 bp of DNA and RNA duplexes to avoid the end effect [18]. For the values of d$L$/d$N$ in Figs. 3 & 6 in the main text, persistence length and stretch modulus shown in Tables S1 and S3, the deviations were calculated based on the values for four equal intervals of MD time in the equilibrium.

**$CoHex^{3+}$ and $Mg^{2+}$ distributions around duplexes: DNA and RNA duplexes**

To describe the binding of $CoHex^{3+}/Mg^{2+}$, we performed the analyses on $CoHex^{3+}/Mg^{2+}$ distributions around DNA and RNA. Specifically, following our previous work [3], we calculated the binding fractions of cations to the major groove, the minor groove, and the phosphates. The cations binding to the grooves were defined as those within the helical radius of 12 Å and in the respective minor and major grooves [3]. The cations binding to phosphates were defined as the cations other than those in grooves and within a radial distance of 6 Å from phosphate atoms. The distributions of $CoHex^{3+}$ and $Mg^{2+}$ around DNA and RNA duplexes were listed in Table S2 in the Supplementary Material.

**Supplementary Figures**

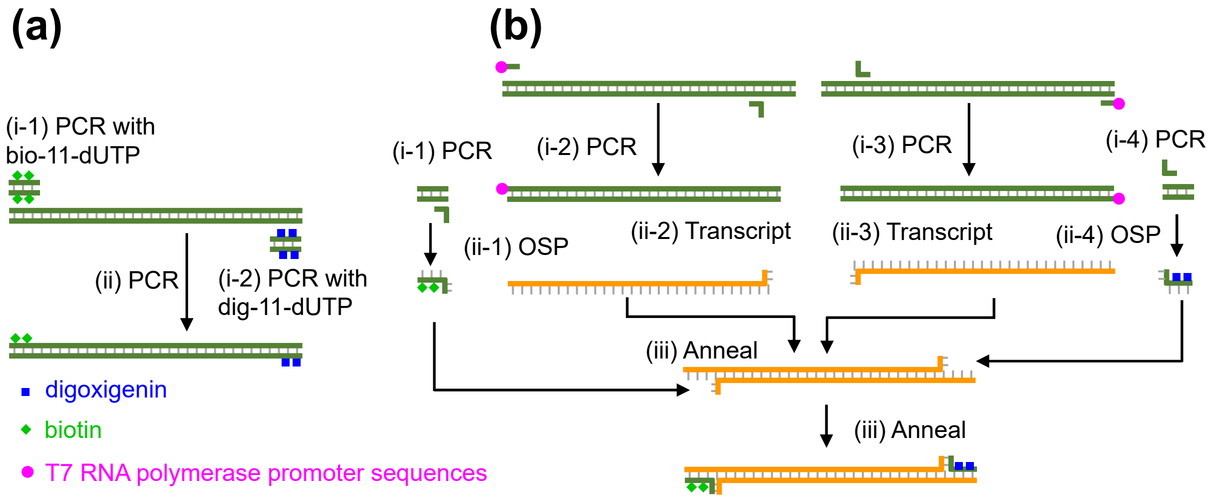


FIG. S1 The preparation of the DNA (a) and RNA (b) constructs.

**(a)** The torsion-constrained DNA construct [1].

(i-1) Make a multiple-biotin-labeled short DNA fragment by PCR using 13K_F and 13K_Rs as primers together with 30% biotin-11-dUTP (Thermo Fisher Scientific) and using lambda DNA as the template.

(i-2) Make a multiple-digoxigenin-labeled short DNA fragment by PCR using 13K_Fs and 13K_R as primers together with 30% digoxigenin-11-dUTP (Roche) and using lambda DNA as the template.

(ii) Make the torsion-constrained DNA construct by PCR using the two labeled short DNA fragments as megaprimers and using lambda DNA as the template.

**(b)** The torsion-constrained RNA construct [2, 3].

(i-1) Make a short DNA fragment through PCR using 13K_F and 13K_Rs as primers and using lambda DNA as the template.

(ii-1) Generate the multiple-biotin-labeled ssDNA through one-sided PCR (OSP) PCR using 13K_ RsL as the primer together with 30% biotin-11-dUTP.

(i-2) Make a long DNA fragment through PCR using 13K_FT7 and 13K_R1L as primers and using the lambda DNA as the template.

(ii-2) Generate an ssRNA strand using T7 RNA polymerase.

(i-3) Make a long DNA fragment through PCR using 13K_F1L and 13K_RT7 as primers and using lambda DNA as the template.

(ii-3) Generate an ssRNA strand using T7 RNA polymerase.

(i-4) Make a short DNA fragment through PCR using 13K_Fs and 13K_R as primers and using lambda DNA as the template.

(ii-4) Generate the multiple-digoxigenin-labeled ssDNA through OSP using 13K_FsL as the primer together with 30% digoxigenin-11-dUTP.

(iii) Anneal above two ssRNA strands and two ssDNA strands (four strands in total) together equimolarly through a temperature process containing a one-hour incubation step at 65 °C followed by a one-hour slow cooling process from 65 °C to 35 °C (-0.5 °C /min).

We synthesized all the DNA oligos in Sangon Biotech (Shanghai) Co., Ltd. with the following sequences.

*13K_F: GCTTGGCTCTGCTAACACGTTGCTCATAGGAG*

*13K_FT7: TAATACGACTCACTATAGGGCTCTGCTAACACGTTGCTCATAGGAG*

*13K_R: AATTTAGCCCTTCAATCGCCAGAGAAATCTAC*

*13K_RT7: TAATACGACTCACTATAGGGAATTTAGCCCTTCAATCGCCAGAGAAATCTAC*

*13K_Rs: CAGCTACAGTCAGAATTTATTGAAGCAA*

*13K_Fs: CCCTAAGACCTTTAATATATCGCCAAATAC*

*13K_FsL:CATGCAATTATTGTGAGCAATACACACGCGCTTCCCCTAAGACCTTTAATATATCGCCA*

*13K_RsL: CATGCAATTATTGTGAGCAATACACACGCGCTTCCAGCTACAGTCAGAATTTATTGAAG*

*13K_R1L: TCATGCAATTATTGTGAGCAATACACACGCGCTTCGCAACAGATATTGAAGGGGAGC*

*13K_F1L: TCATGCAATTATTGTGAGCAATACACACGCGCTTCCTGAAACGTTGCGGTTGAACTAT*

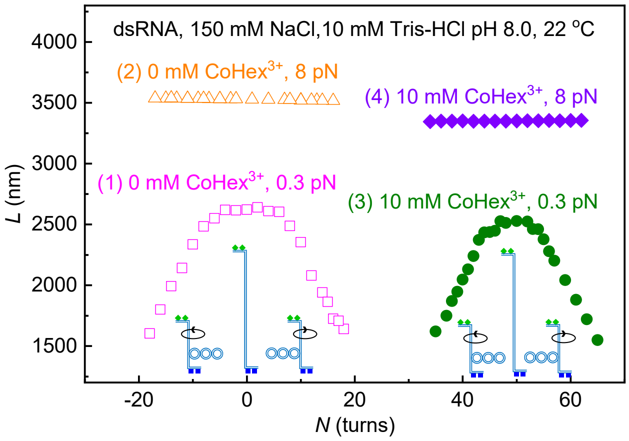


FIG. S2 Twist-stretch coupling measurements. A representative process to measure d*L*/d*N* at different buffer solutions using magnetic tweezers. At each buffer solution, the torsional relaxed point was determined at 0.3 pN. The value of d*L*/d*N* was determined at 8 pN in the range of about ± 20 turns surrounding the torsion relaxed point. (1) In the solution without $CoHex^{3+}$, we rotated the magnets and measured the relationship between the rotations of magnets (*N*) and the extension of RNA (*L*) at 0.3 pN (magenta empty squares). We found the rotation-extension curve demonstrated a bell-shape curve. When overwound or underwound, the RNA duplex shortened due to the formation of plectonemes. We determined the torsion-relaxed point to be the rotation point (*N*=0) where the extension of RNA reached the maximum. (2) Then, we increased the force to 8 pN and measured the rotation-extension curve in the range of about ± 20 turns around the torsion relaxed point (orange empty triangles). We obtained the value of d*L*/d*N* by fitting the rotation-extension curve at 8 pN to a linear function. (3) We rinsed the flow cell with the solution containing 10 mM $CoHex^{3+}$. Then, we measured the bell-shape rotation-extension curve at 0.3 pN to determine the torsion-relaxed point (olive filled cycles). (4) We measured the linear rotation-extension curve at 8 pN and obtained the value of d*L*/d*N* (purple solid diamonds). The magnetic tweezers experiments were performed by following steps:

1. Solution preparation. We purchased 1 M Tris-HCl pH 8.0 stork buffer, Sodium Chloride (NaCl) powder, Magnesium Chloride ($MgCl_2$) powder, Hexaamminecobalt (III) Chloride ($CoHexCl_3$) powder, and Sodium Bicarbonate ($NaHCO_3$) powder from Sigma-Aldrich. We prepared stork solutions of 5 M NaCl, 1 M $MgCl_2$ and 0.2 M $CoHexCl_3$. The final solutions were prepared by diluting the above stork solutions using sterilized DI water. In magnetic tweezers experiments, we rinsed the 50 µL flow cell with at least 2 mL new solution to change the salt concentration.

2. Bead tethering. We anchored the DNA or RNA to the glass slide of the flow cell by a 10-min incubation step in 10 mM Tris-HCl and 500 mM NaCl. Then, we added the microbeads (Dynabeads M-270 Streptavidin) into the flow cell and incubated them for 10 min in 10 mM Tris-HCl and 500 mM NaCl. After that, we passivated the glass and bead surfaces using 10% w/v Methoxy poly(ethylene glycol) succinimidyl valerate (NHS-mPEG, MW 2K, Hunan Huateng Pharmaceutical) in 100 mM $NaHCO_3$ buffer by a two-hour incubation step.

3. Measurements of d*L*/d*N*. In magnetic tweezers experiments, we measured d*L*/d*N* to characterize twist-stretch coupling of DNA and RNA duplexes, where d*N* is the change of twist angle in the unit of helical turn and d*L* is the change of extension of DNA. Fig. S2 shows a representative process we measured d*L*/d*N* at different buffer solutions. The representative process was performed with or without $CoHex^{3+}$ at 150 mM NaCl and 22 °C.

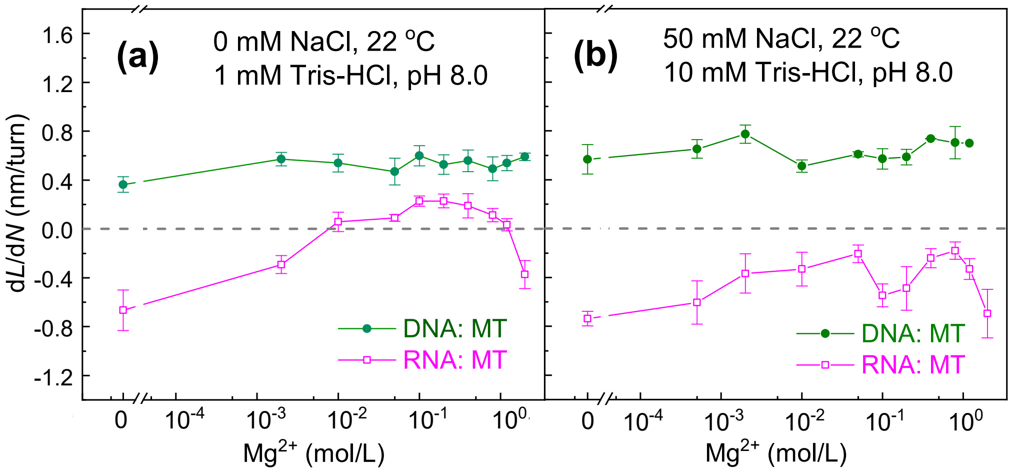


FIG. S3 The twist-stretch coupling parameters (d*L*/d*N*) of DNA and RNA duplexes as functions of $Mg^{2+}$ concentrations at 0 mM NaCl (a) and 50 mM NaCl (b). The experimental coupling parameters (d*L*/d*N*) obtained from more than four molecules are plotted as data points and error bars.

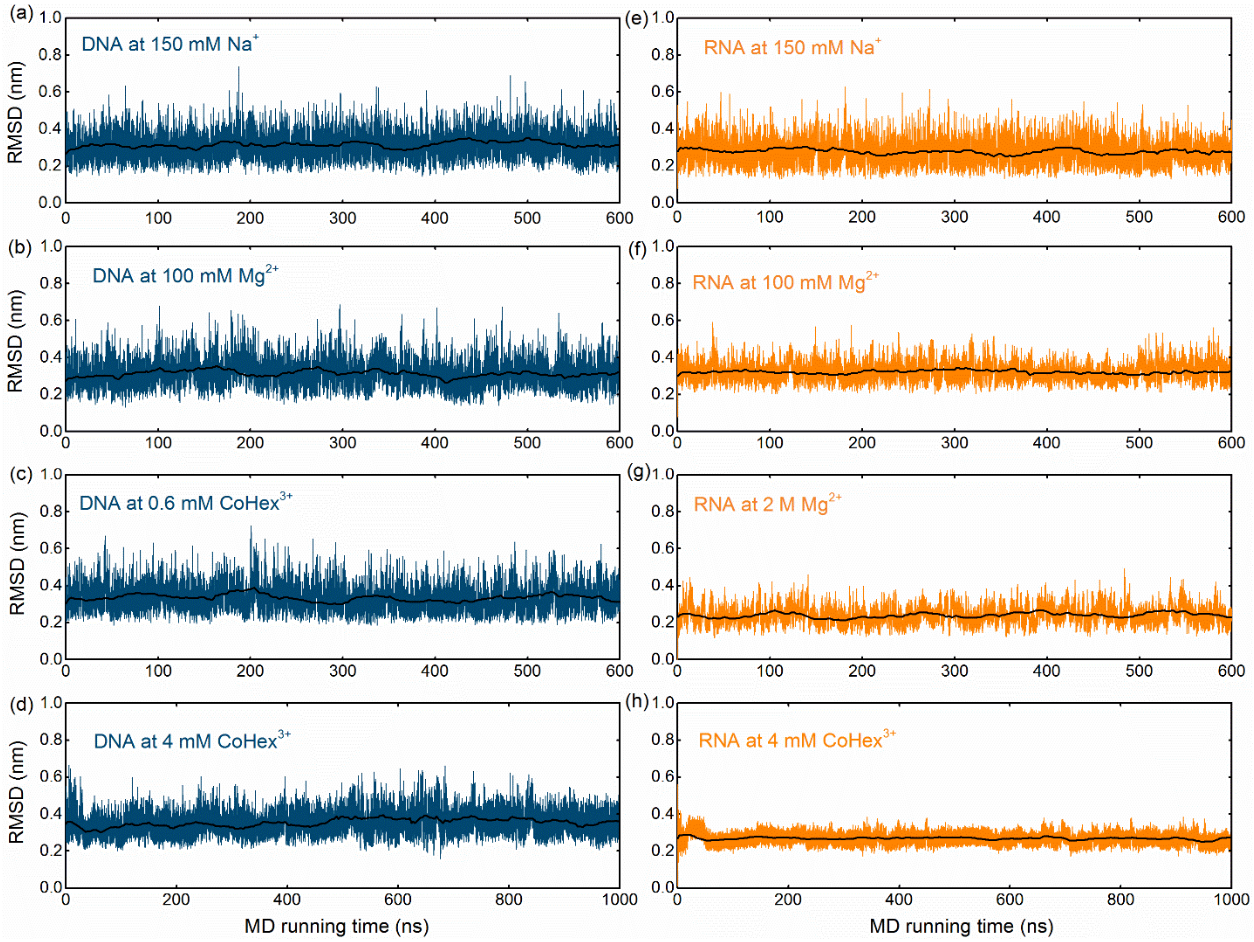


FIG. S4 The root-mean-square deviation (RMSD) versus MD running time for the central 14-bp segment of the DNA (a-d) and RNA (e-h) duplexes. The black lines represent the RMSD values averaged over every 2 ns.

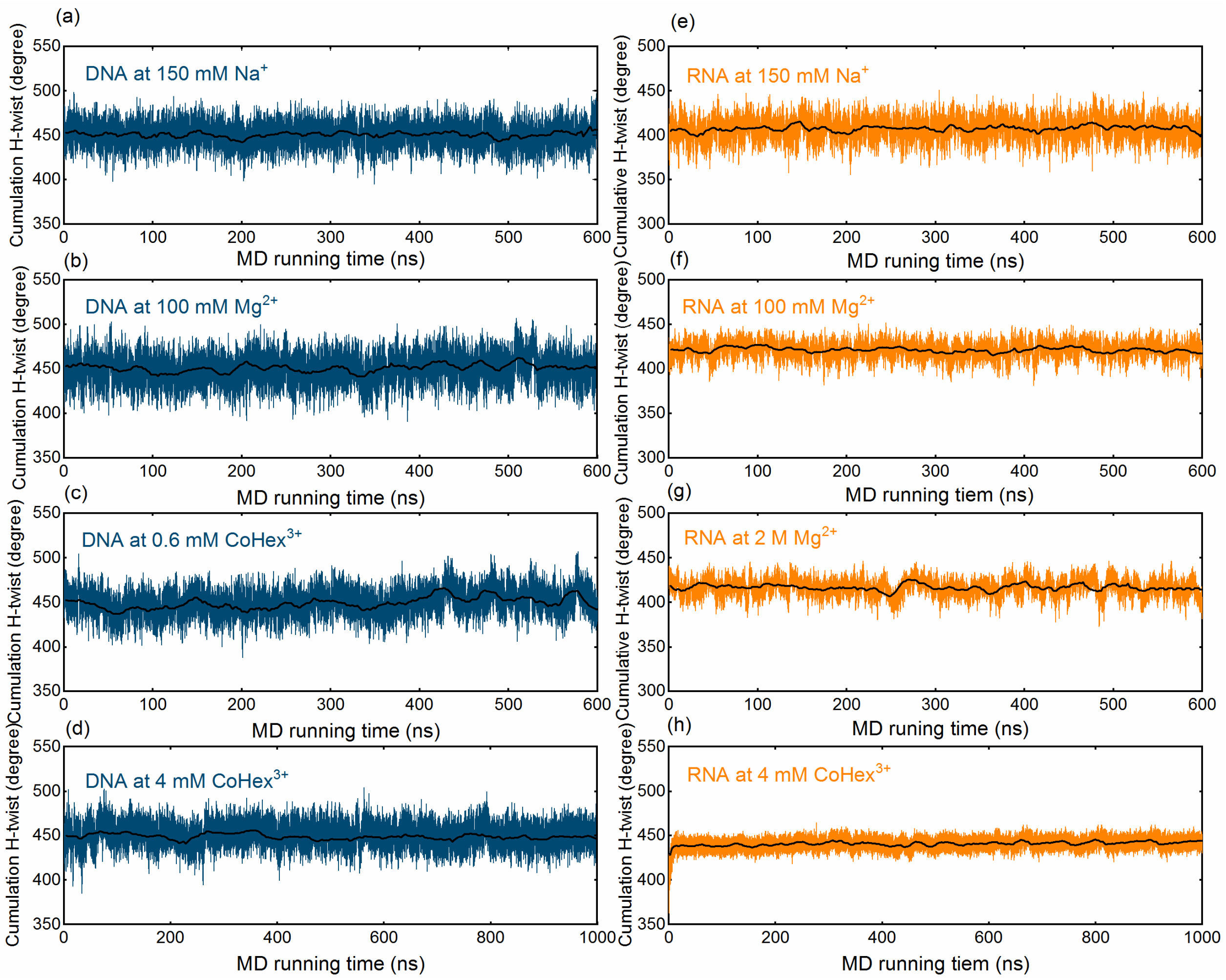


FIG. S5 The cumulative H-twist versus MD running time for the central 14-bp segments of the DNA (a-d) and RNA (e-h) duplexes. The central black lines represent the contour length averaged over every 2 ns.

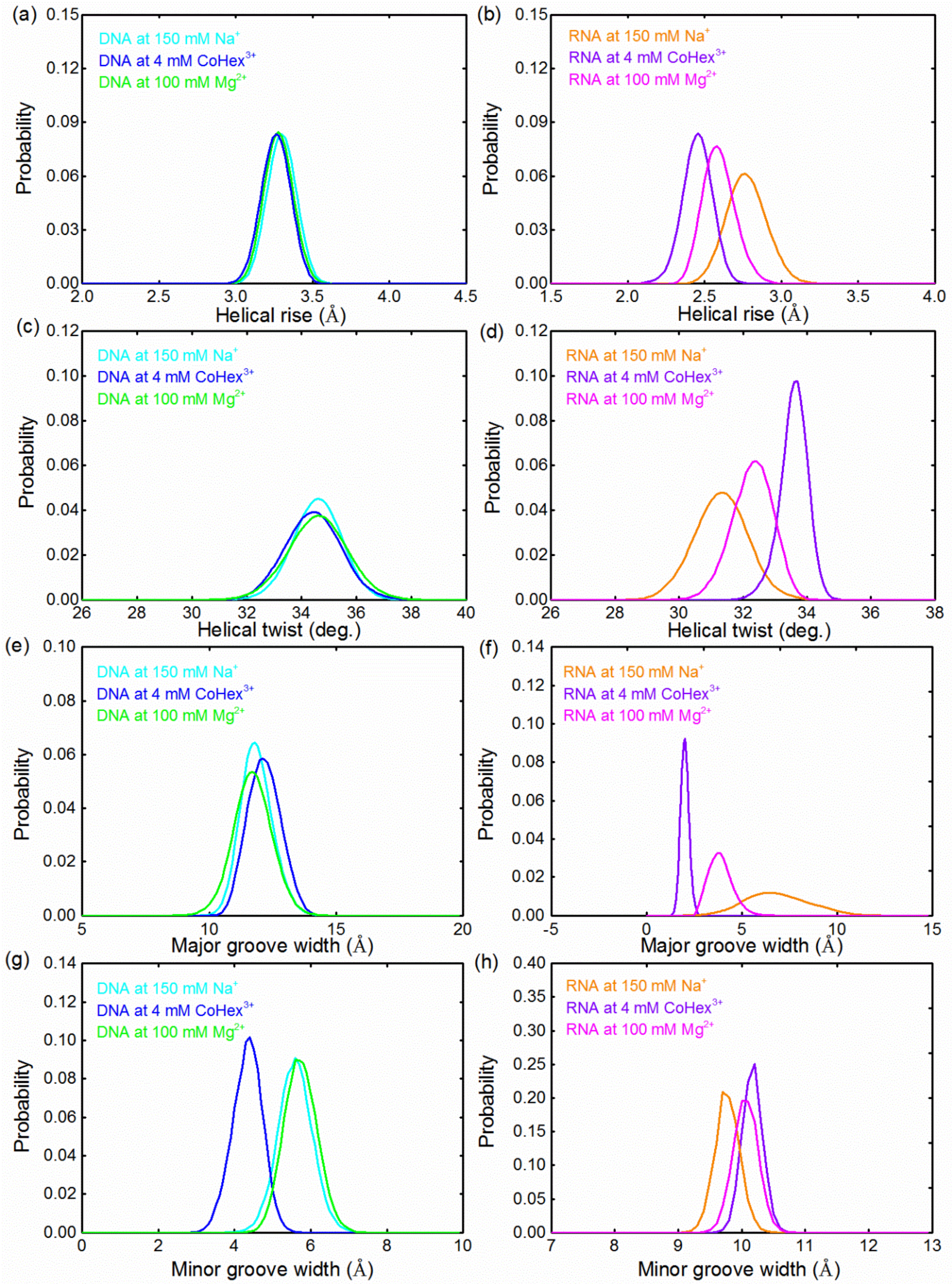


FIG. S6 The probability distributions of helical rise (a, b), helical twist (c, d), major groove width (e, f) and minor groove width (g, h) for the central 14-bp segments of the DNA and RNA duplexes.

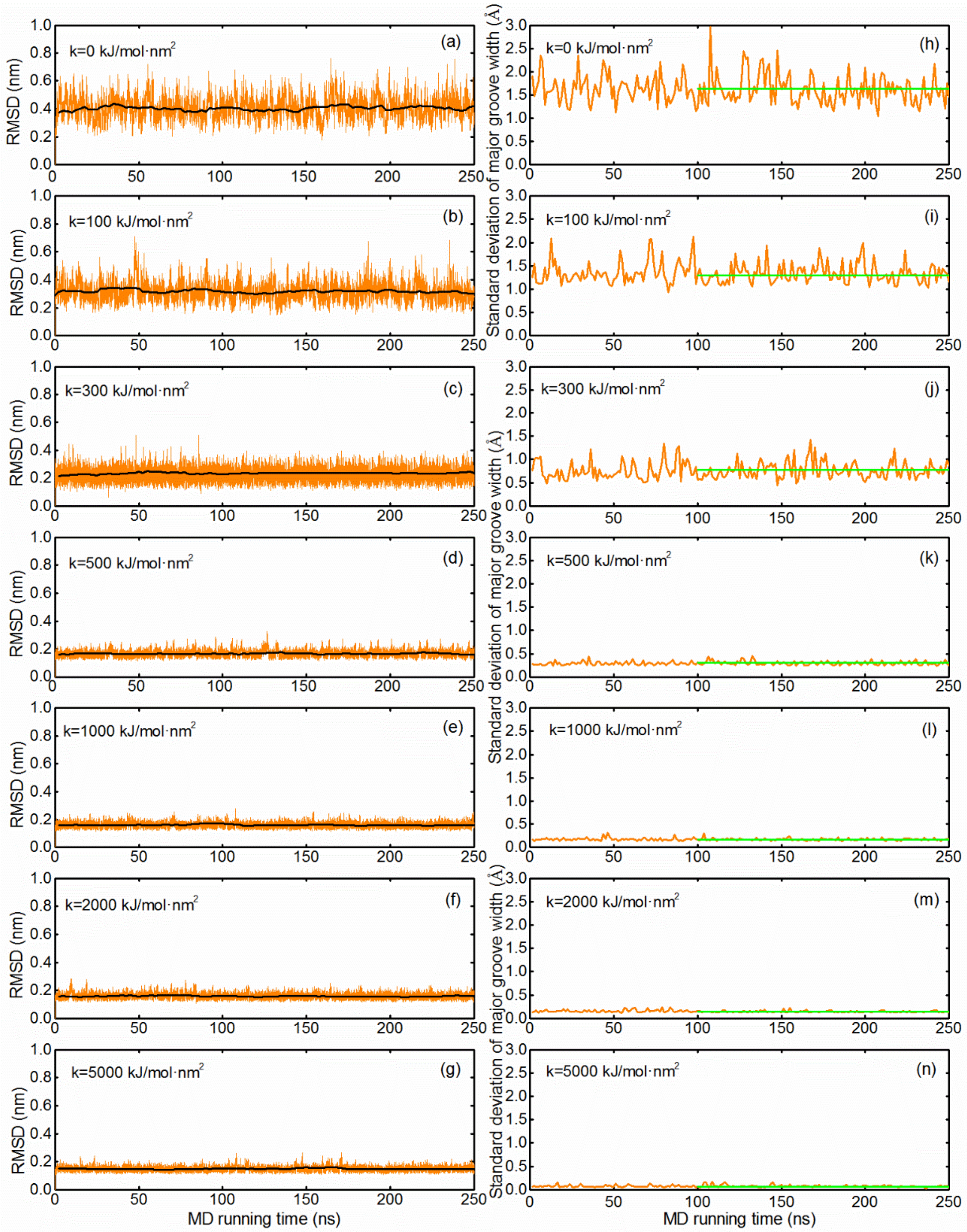


FIG. S7 The root-mean-square deviation (RMSD) from the constrained all-atom MD simulations versus MD running time for the central 14-bp segment of the RNA (a-g) duplex, and the black lines represent the RMSD values averaged over every 2 ns. (h-n) The standard deviations of major groove widths from the constrained all-atom MD simulations versus MD time for the central 14-bp segments of the RNA duplex, which are calculated over every 2 ns with respect to the mean groove widths in the range of 100-250 ns (denoted by straight green lines).

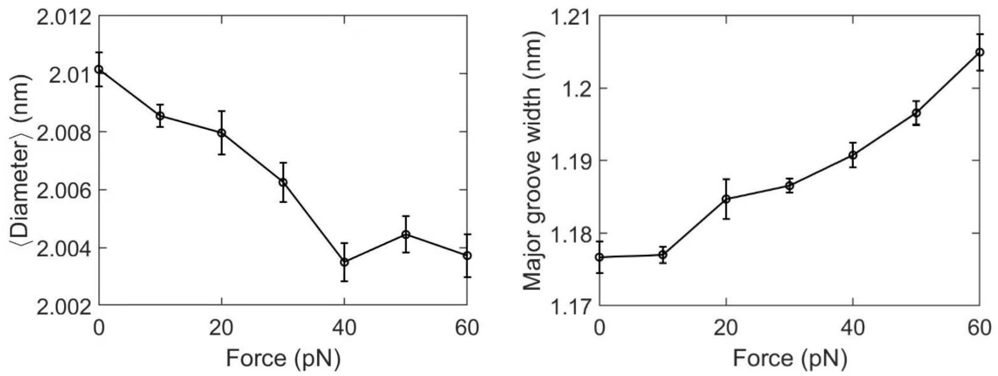


FIG. S8 The mean diameter (left panel) and major groove width (right panel) as a function of external stretching force for the DNA duplex from the all-atom MD simulations under stretching force. It appears that the DNA diameter reaches its minimum when the external stretching force is about 40 pN.

### Supplementary Tables

**Table S1:** Microscopic structural parameters for DNA and RNA duplexes.

| | DNA at 150 mM $Na^+$ | DNA at 4 mM $CoHex^{3+}$ [a] | DNA at 10 mM $Na^+$ | DNA at 100 mM $Mg^{2+}$ [b] | DNA at 0.6 mMCoHex$^{3+}$[a] |
|---|---|---|---|---|---|
| Helical rise (Å) | 3.33±0.09 | 3.31±0.09 | 3.31±0.08 | 3.29±0.09 | 3.32±0.91 |
| Helical twist (degree) | 34.62±1.10 | 34.31±1.22 | 34.69±0.91 | 34.56±1.11 | 34.92±0.97 |
| Inclination (degree) | 3.43±2.40 | 2.43±2.49 | 2.78±2.30 | 3.13±2.28 | 2.73±2.43 |
| Helical radius (Å) | 10.14±0.24 | 10.23±0.32 | 10.13±0.22 | 10.12±0.20 | 10.19±0.29 |
| Major groove width (Å) | 11.92±0.64 | 12.35±0.72 | 11.96±0.63 | 11.76±0.77 | 12.36±0.71 |
| Minor groove width (Å) | 5.64±0.46 | 4.34±0.47 | 5.61±0.44 | 5.78±0.44 | 4.64±0.51 |
| d*L*/d*N* (MT) | 0.42 | 0.85[c] | 0.47 | 0.60 | 0.96[c] |
| d*L*/d*N* (MD) | 0.49 | 0.61 | 0.30 | 0.58 | 0.89 |
| | RNA at 150 mM $Na^+$ | RNA at 4 mM $CoHex^{3+}$ [a] | RNA at 10 mM $Na^+$ | RNA at 100 mM $Mg^{2+}$ [b] | RNA at 2 M $Mg^{2+}$ [b] |
| Helical rise (Å) | 2.79±0.13 | 2.46±0.10 | 2.86±0.13 | 2.59±0.10 | 2.73±0.12 |
| Helical twist (degree) | 31.36±0.94 | 33.69±0.48 | 30.68±0.93 | 32.36±0.64 | 32.04±0.87 |
| Inclination (degree) | 15.23±2.72 | 21.62±1.23 | 12.03±2.56 | 19.30±1.93 | 15.81±2.42 |
| Helical radius (Å) | 9.97±0.24 | 9.33±0.16 | 10.11±0.26 | 9.67±0.22 | 9.79±0.23 |
| Major groove width (Å) | 6.90±1.66 | 2.04±0.22 | 7.21±1.68 | 3.84±1.13 | 6.01±1.34 |
| Minor groove width (Å) | 9.83±0.19 | 10.21±0.16 | 9.62±0.21 | 10.15±0.18 | 9.87±0.18 |
| d*L*/d*N* (MT) | -0.58 | 0.40[c] | -0.65 | 0.23 | -0.37[c] |
| d*L*/d*N* (MD) | -0.76 | 0.71 | -0.79 | 0.25 | -0.49 |

[a]In 150 mM $Na^+$ buffer.
[b]In 10 mM $Na^+$ buffer.
[c]The data obtained from the linear interpolation for the experimental data.

**Table S2:** Distributions of $CoHex^{3+}$ and $Mg^{2+}$ around DNA and RNA duplexes revealed by MD simulations.

| | phosphate | Major groove | Minor groove |
|---|---|---|---|
| 4 mM $CoHex^{3+}$ on DNA | 59% | 24% | 17% |
| 100 mM $Mg^{2+}$ on DNA | 36% | 43% | 20% |
| 0.6 mM $CoHex^{3+}$ on DNA | 51% | 30% | 19% |
| 4 mM $CoHex^{3+}$ on RNA | 15% | 75% | 10% |
| 100 mM $Mg^{2+}$ on RNA | 21% | 62% | 17% |
| 2 M $Mg^{2+}$ on RNA | 51% | 40% | 9% |

**Table S3:** Microscopic structural parameters for RNA duplex with different constraint constants *k* across major groove from the constrained MD simulations.

| kJ/mol·nm$^2$ | *k*=0 | *k*=100 | *k*=300 | *k*=500 | *k*=1000 | *k*=2000 | *k*=5000 |
|---|---|---|---|---|---|---|---|
| Helical rise (Å) | 2.81±0.13 | 2.72±0.11 | 2.62±0.10 | 2.47±0.09 | 2.46±0.08 | 2.46±0.05 | 2.45±0.04 |
| Helical twist (degree) | 31.33±0.90 | 31.93±0.78 | 32.81±0.66 | 33.64±0.42 | 33.74±0.38 | 33.74±0.36 | 34.01±0.35 |
| Inclination (degree) | 14.20±2.58 | 18.73±2.10 | 20.64±1.48 | 22.39±1.36 | 22.51±1.21 | 22.53±1.20 | 22.71±1.16 |
| Helical radius (Å) | 9.98±0.26 | 9.89±0.22 | 9.62±0.19 | 9.45±0.12 | 9.46±0.03 | 9.41±0.03 | 9.32±0.02 |
| Persistence length (nm) | 54.30±0.5 | 56.50±0.4 | 77.80±0.3 | 107.80±0.3 | 131.70±0.2 | 151.50±0.2 | 167.90±0.1 |
| Strength module (pN) | 496±65 | 673±58 | 780±53 | 887±41 | 1149±32 | 2685±27 | 4033±21 |
| Major groove width (Å) | 7.13±1.63 | 5.97±1.31 | 3.65±0.78 | 2.05±0.29 | 2.04±0.17 | 2.01±0.14 | 1.99±0.07 |
| Minor groove width (Å) | 9.78±0.20 | 9.84±0.18 | 10.03±0.17 | 10.25±0.17 | 10.25±0.17 | 10.27±0.16 | 10.34±0.15 |

**References**

[1] D.H. Paik, V.A. Roskens and T.T. Perkins, Nucleic Acids Res. **41**, 19 (2013).
[2] Y.-J. Yang, L. Song, X.-C. Zhao, C. Zhang, W.-Q. Wu, H.-J. You, H. Fu, E.-C. Zhou and X.-H. Zhang, ACS synthetic biology. **8**, 7 (2019).
[3] H. Fu, C. Zhang, X.-W. Qiang, Y.-J. Yang, L. Dai, Z.-J. Tan and X.-H. Zhang, Phys. Rev. Lett. **124**, 5 (2020).
[4] R.S. Mathew-Fenn, R. Das and P.A. Harbury, Science. **322**, 5900 (2008).
[5] A. Pérez, I. Marchán, D. Svozil, J. Sponer, T.E. Cheatham III, C.A. Laughton and M. Orozco, Biophys. J. **92**, 11 (2007).
[6] I.S. Joung and T.E. Cheatham III, The Journal of Physical Chemistry B. **112**, 30 (2008).
[7] T. Sun, A. Mirzoev, N. Korolev, A.P. Lyubartsev and L. Nordenskiöld, The Journal of Physical Chemistry B. **121**, 33 (2017).
[8] J. Lipfert, G. Skinner, J. Keegstra, T. Hensgens, T. Jager, D. Dulin, M. Köber, Z. Yu, S. Donkers, F. Chou, et al., Proc. Natl. Acad. Sci. U.S.A. **111**, 43 (2014).
[9] Y.-Y. Wu, Z.-L. Zhang, J.-S. Zhang, X.-L. Zhu and Z.-J. Tan, Nucleic Acids Res. **43**, 12 (2015).
[10] Z.-J. Tan and S.-J. Chen, Biophys. J. **99**, 5 (2010).
[11] K. Xi, F.-H. Wang, G. Xiong, Z.-L. Zhang and Z.-J. Tan, Biophys. J. **114**, 8 (2018).
[12] D.M. York, T.A. Darden and L.G. Pedersen, The Journal of Chemical Physics. **99**, 10 (1993).
[13] S. Miyamoto and P.A. Kollman, J. Comput. Chem. **13**, 8 (1992).
[14] E.D. Merkley, S. Rysavy, A. Kahraman, R.P. Hafen, V. Daggett and J.N. Adkins, Protein Sci. **23**, 6 (2014).
[15] A.E. Torda, R.M. Scheek and W.F. Van Gunsteren, Chem. Phys. Lett. **157**, 4 (1989).
[16] R. Lavery, M. Moakher, J.H. Maddocks, D. Petkeviciute and K. Zakrzewska, Nucleic Acids Res. **37**, 17 (2009).
[17] L. Bao, X. Zhang, Y.-Z. Shi, Y.-Y. Wu and Z.-J. Tan, Biophys. J. **112**, 6 (2017).
[18] Y.-Y. Wu, L. Bao, X. Zhang and Z.-J. Tan, J. Chem. Phys. **142**, 12 (2015).
[19] I. Faustino, A. Pérez and M. Orozco, Biophys. J. **99**, 6 (2010).
[20] A. Noy and R. Golestanian, Phys. Rev. Lett. **109**, 22 (2012).
